# Importance of cysteines for the binding of S100A6 to the RAGE receptor – Towards a first molecular model for S100 covalent dimerization

**DOI:** 10.1101/2025.07.25.666771

**Authors:** Maria Demou, Laure Yatime

**Affiliations:** Laboratory of Pathogens and Host Immunity (LPHI), University of Montpellier, INSERM UA15, CNRS UMR5294, Montpellier, France

**Keywords:** S100 proteins, RAGE receptor, cysteines, disulfide-crosslinking, homodimerization

## Abstract

Extracellular S100 proteins act as alarmins and trigger pro-inflammatory signaling cascades by activating their cognate cell-surface receptor RAGE, thereby contributing to both normal and pathological inflammation depending on the physiological context. These ligand-receptor interactions occur in an oxidative environment that is known to induce post-translational modifications, notably on the cysteine residues present in S100 proteins. How cysteine oxidation affects the architecture of S100 proteins and their interaction with RAGE remains poorly understood as most *in vitro* studies employ cysteine mutants or reduced conditions. Using our model protein S100A6 and size exclusion chromatography-based binding assays in non-reducing conditions, we here demonstrate that the unique cysteine of S100A6, Cys3, is essential for the binding to RAGE. We further show that full complexation can be restored by introducing a cysteine at conserved position 84, where a Cys residue is found in at least ten other RAGE-binding S100 proteins. Structural analysis of the resulting complex between RAGE ectodomain and S100A6 mutant Y84C further reveals that the presence of Cys84 induces the formation of a covalent disulfide bond between the two S100A6 protomers, thus stabilizing the same RAGE-bound S100A6 conformation as with the WT protein. Finally, modeling of other S100 proteins in the RAGE-bound conformation suggests that this covalent S100 dimer architecture may be adopted by other members of the family, already reported to form disulfide-crosslinked oligomeric species. Altogether, our findings highlight the importance of S100 cysteines for the binding to RAGE and provide a first molecular model for S100 covalent homodimerization.

## Introduction

S100 proteins are small calcium-binding proteins from the EF-hand superfamily. They are found almost exclusively in vertebrates, where they exert pleiotropic functions, both intra- and extracellularly [1]. Inside cells, they regulate calcium-dependent processes linked to calcium homeostasis, metabolism, cell proliferation, motility or cytoskeleton rearrangement [2]. Outside cells, they act as immune-activating alarmins [3] by binding to cognate cell-surface receptors, such as the receptor for advanced glycation end-products (RAGE), through which they trigger downstream pro-inflammatory signaling cascades [4,5]. S100 proteins also take part in the antimicrobial response, thanks to their ability to sequester metal ions with high affinity, a property that leads to nutritional immunity [6]. Their functional versatility is thought to arise from their propensity to form distinct assemblies that may exert distinct biological functions [7]. Hence, S100 proteins all form dimers as the minimal, functional unit; they can also arrange into higher order oligomers, such as tetramers, hexamers or octamers [8–11]. Furthermore, they can form heterocomplexes, like the highly proinflammatory S100A8/A9 heterodimer. Most importantly, their tertiary and quaternary architecture can also be modulated in response to binding of divalent cations or following post-translational modifications such as phosphorylation, nitrosylation or oxidation [12–15].

The impact of redox conditions on the structure-function relationships of S100 proteins is still lacking in-depth investigation, despite the fact that these proteins undergo a drastic environmental change upon their relocalization from the reduced intracellular compartment to the highly oxidative extracellular space. Interestingly, a few reports have shown that oxidation of S100 proteins can lead to the formation of disulfide-crosslinked species, both *in vitro* and *in vivo*, and that such process serves as a regulatory switch for their alarmin function. For example, oxidation of S100A8/A9 leads to the formation of a covalent heterodimer through the linkage of Cys3 of S100A9 with Cys42 of S100A8. This covalent form shows protective and pro-resolutive properties as compared to the reduced form which sustains high inflammation [16,17], mostly due to a higher sensitivity to proteolytic degradation [18]. Oxidized S100A4 also assembles into disulfide-crosslinked oligomers that fail to interact with protein phosphatase 5 (PP5) and further prevent PP5 activation by S100A1, thereby modulating apoptosis under oxidative stress conditions [19]. Of importance, the interaction of S100 proteins with RAGE may also be modulated by cysteine-dependent oxidative processes. Hence, oxidation of S100B generates disulfide-crosslinked octamers that display enhanced binding to RAGE and can induce RAGE-dependent secretion of vascular endothelial growth factor (VEGF), unlike non-covalent S100B dimers [15].

These examples highlight the importance of cysteine residues for the extracellular functions of S100 proteins, including for their interaction with effectors like RAGE. How cysteine-dependent oxidative modifications affect the quaternary architecture of S100 proteins at the fine molecular level and how cysteines may modulate S100’s interaction with RAGE remains however unclear, notably because many *in vitro* studies are performed with recombinant S100 proteins purified under reducing conditions or for which cysteines have been mutated away [20–22]. Here, we address this question by focusing on the model S100 protein studied in our group, S100A6 [23]. Using a combination of size exclusion chromatography (SEC) and X-ray crystallography experiments, we demonstrate that the presence of a cysteine at two critical conserved positions within the S100 protein family is pivotal in enabling S100A6 binding to the RAGE ectodomain. We further propose a first atomic model describing the quaternary organization of a disulfide-crosslinked S100 dimer, rationalized from a mutated form of S100A6. Finally, based on computational modeling, we discuss the possibility that this architecture may be adopted by other RAGE-binding S100 proteins known to form covalent oligomers.

## Results

### Cys3 is essential for the binding of S100A6 to RAGE

S100A6 bears a single cysteine at position 3 that promotes formation of S100A6 disulfide-crosslinked dimers [24]. Interestingly, oxidative modification of S100A6 was shown to impair its interaction with effectors like PP5 [25]. The importance of Cys3 or Cys3-dependent structural modification of S100A6 on its binding to RAGE has however not been addressed yet. To investigate the role of this oxidation-sensitive residue, we monitored the formation of RAGE:S100A6 complexes in solution, using SEC. As shown in Figure 1B (dark blue curve), incubation of the RAGE VC1C2 fragment (full-length ectodomain) with WT S100A6, in the presence of calcium, allows the formation of a novel species eluting earlier than each isolated protein (Fig.1B, red and orange curves for isolated VC1C2 and S100A6, respectively). This species contains both VC1C2 and S100A6 as visualized by SDS-PAGE (Fig.1F, lane 2), thus revealing the formation of a stable complex between the two proteins. Formation of this RAGE:S100A6 complex is strictly calcium-dependent since no earlier peak is detected when the two proteins are incubated in the absence of calcium (Fig.1B, light blue curve).

**Figure 1.**
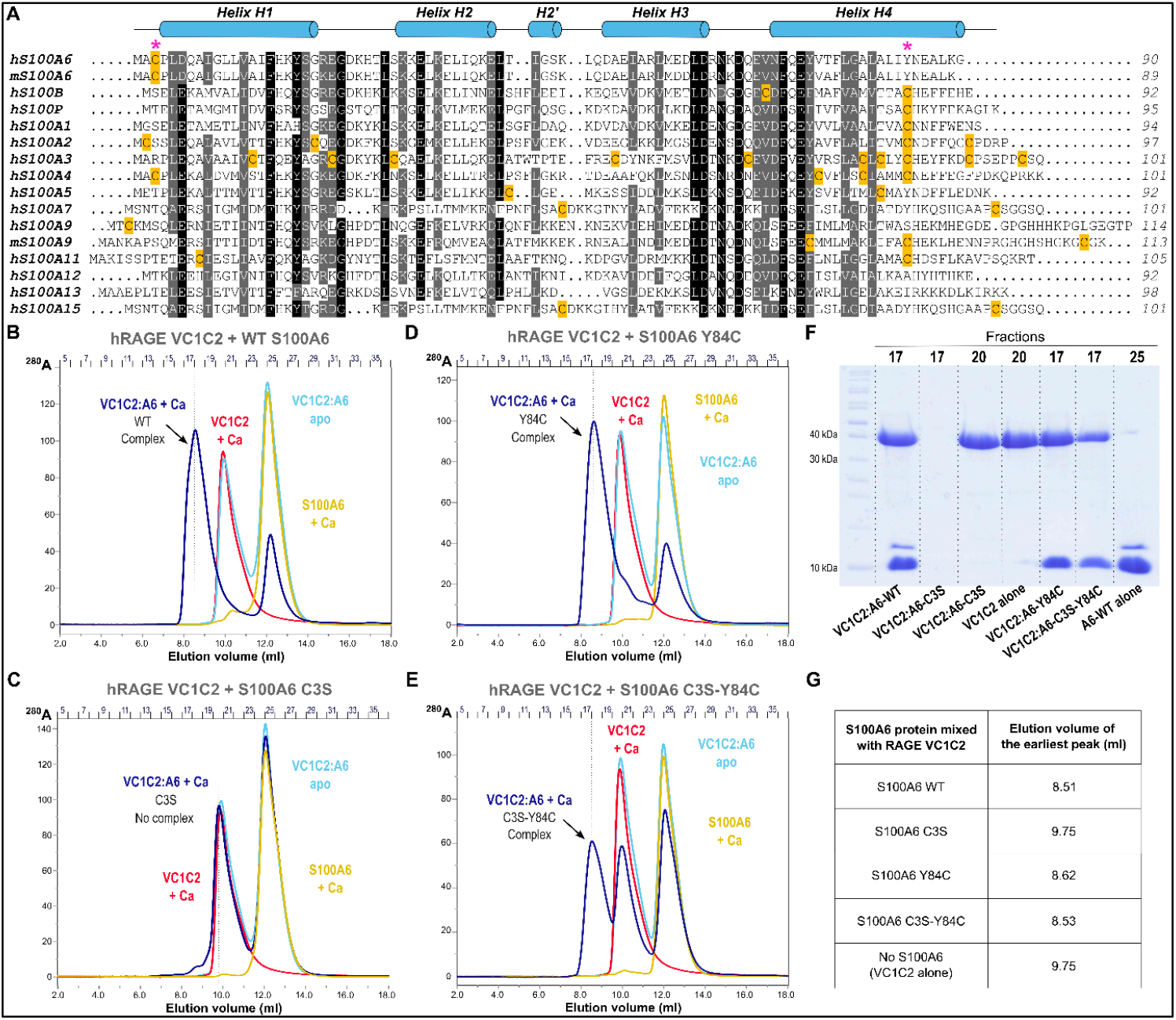
Interaction of S100A6 with the RAGE ectodomain is dependent on cysteine residues. **A**. Sequence alignment of the human S100 proteins known to bind to the RAGE receptor. Murine S100A6 and S100A9 are also included in the alignment. Sequences are colored according to sequence conservation. The secondary structure elements of S100 proteins (α-helices) are indicated above the alignment, based on S100A6 tridimensional structure. Cysteine residues are highlighted in orange. The 2 positions for the conserved Cys residues investigated in this study (Cys3 and Cys84) are marked by pink asterisks above the alignment. **B. to E**. Formation of RAGE:S100A6 complexes as assessed by analytical SEC with either WT S100A6 (panel B) or S100A6 mutants C3S (panel C), Y84C (panel D) and C3S-Y84C (panel E). All SEC experiments were performed on a 24 ml Superdex 75 Increase column (Cytiva) equilibrated in 20 mM Tris-HCl pH 7.5, 200 mM NaCl. Elution profiles corresponding to the 1:10 molar ratio mixes between RAGE and S100A6 are shown in dark blue (incubation in the presence of 5 mM CaCl_2_) or light blue (incubation without Ca). Elution profiles for the isolated proteins are shown in red (RAGE VC1C2) or orange (S100A6). **F**. SDS-PAGE analysis of selected fractions from the different elution runs. The number of the analyzed fractions is indicated above the gel and the corresponding chromatogram is indicated below the gel, for lanes 2 to 8. Lane 1 corresponds to the molecular weight marker (PageRuler™ unstained protein ladder, Thermo Fisher). **G**. Elution volume of the earliest peak observed for the different RAGE:S100A6 mixes incubated in the presence of calcium (dark blue chromatograms). The elution volume of the peak corresponding to RAGE alone (red curve) is indicated for comparison.

Next, we investigated complex formation with the C3S mutant of S100A6. As shown in Figure 1C (dark blue curve), no complex is formed when RAGE-VC1C2 is incubated with S100A6-C3S. The chromatogram obtained for the mix in the presence of calcium perfectly superimposes with that obtained for the mix in absence of calcium (Fig.1C, light blue curve) or with the combined chromatograms of the two isolated proteins (Fig.1C, red and orange curves). SDS-PAGE analysis confirms that the fractions where the WT RAGE:S100A6 eluted previously do not contain any protein (Fig.1F, lane 3), while the earliest peak observed for the VC1C2:S100A6-C3S mix, eluting at the same volume as isolated VC1C2 domain (Fig. 1G), only contains VC1C2 (Fig.1F, lane 4). Thus, in solution, S100A6 is unable to interact with RAGE full-length ectodomain when Cys3 is mutated away.

### Replacement of Cys3 by Cys84 restores complex formation between S100A6 and RAGE

This finding prompted us to investigate whether a cysteine was present at the beginning of helix H1 in other RAGE-binding S100 proteins, which could hint for a more universal importance of Cys residues for RAGE:S100 complex formation. Sequence alignment of the S100 proteins known to interact with RAGE [26] revealed that only few S100 proteins possess a cysteine near their N-terminus, namely S100A2, S100A4 and human S100A9 (but not murine S100A9) (Fig. 1A). In contrast, many RAGE-interacting S100 proteins bear a Cys residue in the second half of their last helix (helix H4). In particular, a highly conserved Cys is found at the position equivalent to Tyr84 in S100A6. To evaluate the importance of having a cysteine at this position, we introduced the Y84C mutation in S100A6, both in WT and C3S background.

S100A6-Y84C behaves similarly as the WT protein (Fig. 1D; Fig. 1F, lane 6), also forming a Ca^2+^-dependent complex with RAGE that elutes at a volume of 8.6 mL (Fig. 1G), likewise the WT complex. Most interestingly, while the C3S mutation totally abrogates complex formation, the S100A6 C3S-Y84C double mutant regains ability to interact with RAGE, the presence of the two proteins in the peak eluting at 8.5 mL being confirmed by SDS-PAGE (Fig.1F, lane 7). This complex is specific since it is only obtained in the presence of calcium, proving that it is not due to aberrant Cys-Cys crosslinking between RAGE and S100A6. Full complexation is however not reached as compared to the WT proteins, which may indicate a slower rate of association or a less stable complex. Nevertheless, these data show that replacement of Cys3 by Cys84 allows to restore S100A6 interaction with RAGE ectodomain.

### Presence of a cysteine at position 84 allows to form a disulfide-crosslinked S100A6 dimer

To gain insights at the molecular level on how the presence of a cysteine at position 84 may promote RAGE:S100A6 complexation similarly as when cysteine 3 is present, we sought to crystallize the mutated complexes. We could only obtain crystals for the VC1C2:S100A6-Y84C complex that diffracted to 2.35 Å resolution (Table I). Phasing was achieved using molecular replacement, with the structure of the WT complex as a search template (PDB_ID 4P2Y; [23]). The refined atomic model is displayed in Figure 2A and reveals a 2:2 complex that is highly similar to the complex obtained with WT S100A6, with an overall root-mean-square deviation (RMSD) on Cα atoms of 0.4 Å between the WT and mutant structures.

**Table I.**
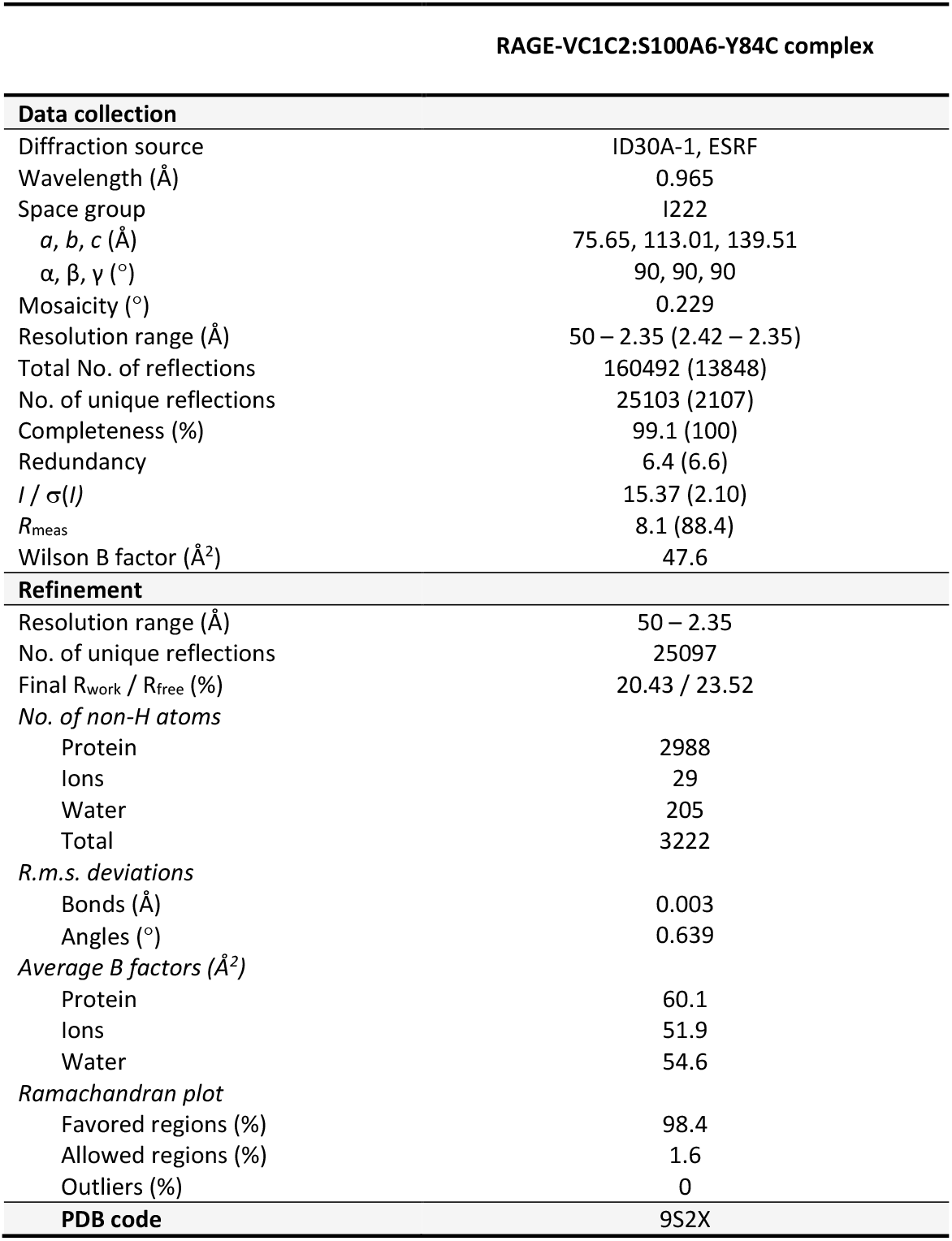
Data collection and refinement statistics. Values indicated in parentheses correspond to the last shell of resolution.

**Figure 2.**
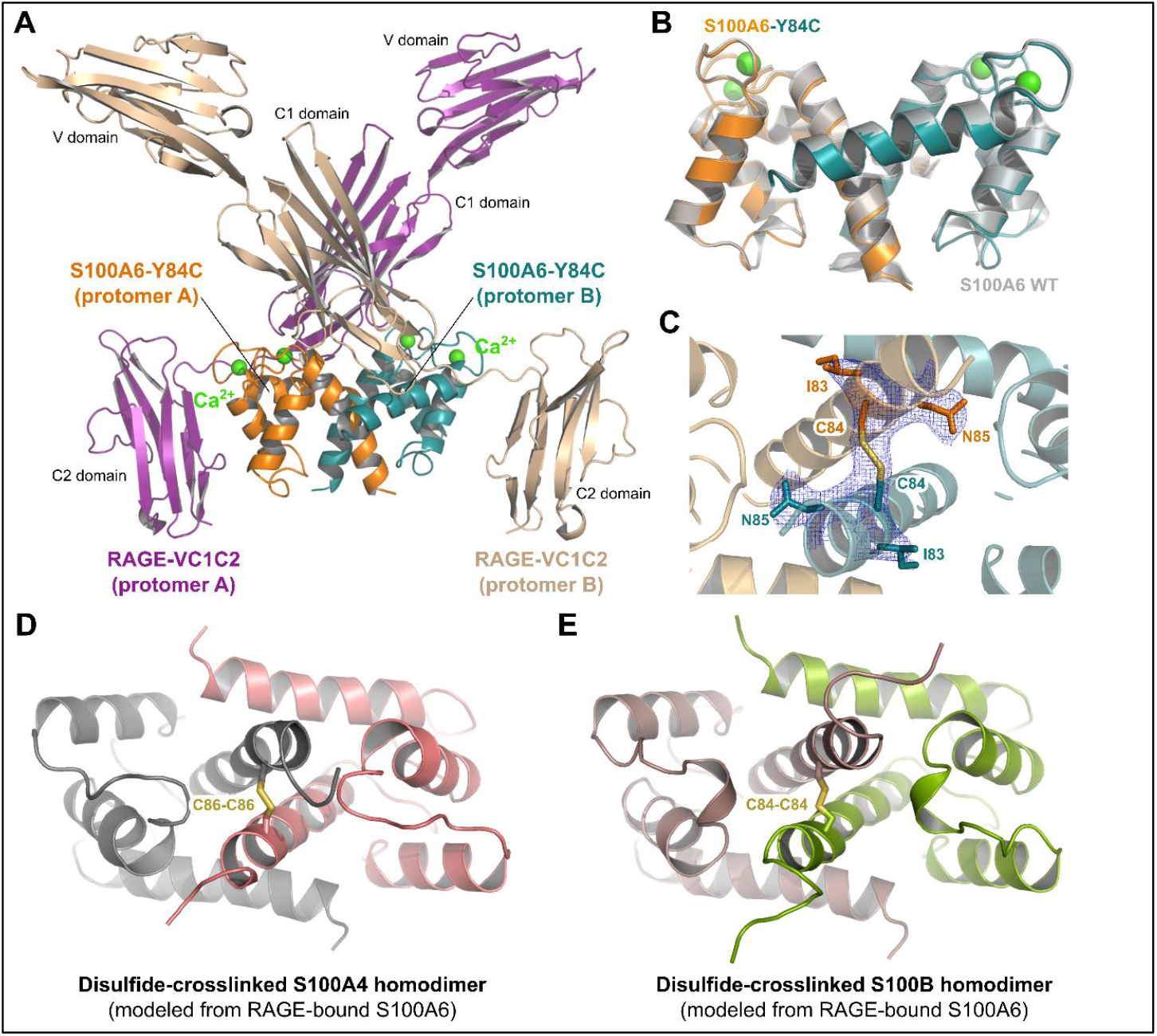
A first putative model for S100 disulfide-crosslinked homodimers. **A**. Crystallographic structure of the RAGE:S100A6-Y84C complex determined at 2.35 Å resolution. The biological assembly is similar to that of the WT complex, i.e. it encompasses two molecules of RAGE bound to one S100A6 homodimer (2:2 complex). **B**. Superimposition of the RAGE-bound S100A6-Y84C homodimer (in orange and blue) with the RAGE-bound WT S100A6 (in gray; PDB_ID 4P2Y [23]). **C**. Zoom on the C-terminal helices (H4) of both S100 protomers in the RAGE-bound S100A6-Y84C homodimer. The two protomers are covalently linked via an intermolecular disulfide bridge (Cys84-Cys84). The single SA-omit map calculated in PHENIX.REFINE after removing S100A6 residues Ile83 to Asn85 in the model is displayed as blue mesh around this region (contour at 1σ), allowing to visualize the clearly defined density for the SS-bond. **D**. Modeling of S100A4 homodimer in the RAGE-bound S100A6 conformation highlights the possible formation of an intermolecular disulfide crosslink between the Cys86 from the two protomers. **E**. Modeling of S100B homodimer in the RAGE-bound S100A6 conformation highlights the possible formation of an intermolecular disulfide crosslink between the Cys84 from the two protomers. The models were built manually by superimposing the structure of the S100 protomers from Ca^2+^-bound human S100A4 (PDB_ID 2Q91 [39]) or Ca^2+^-bound human S100B (PDB_ID 2H61 [10]) onto the two protomers of the RAGE-bound S100A6-Y84C homodimer (this study).

In the mutant complex, S100A6-Y84C adopts the same homodimeric conformation as that of RAGE-bound WT S100A6 (Fig. 2B), i.e. helices H1 and H4 from the first S100A6 protomer are arranged in an antiparallel manner with the same helices from the second S100A6 protomer. Most interestingly, when zooming on the region of the atomic model near position 84 within S100A6, we observe that the two mutated Cys84 from each protomer are in such close proximity that a disulfide bridge is formed between them, thus giving rise to a disulfide-crosslinked S100A6-Y84C dimer (Fig. 2C). The presence of this S-S linkage is not due to a modeling bias since it is visible in the single SA-omit map calculated from a model where residues 83 to 85 of S100A6 have been omitted (Fig. 2C, blue mesh). Our data thus suggest that introduction of a cysteine at position 84 in S100A6 stabilizes the formation of the RAGE-bound S100A6 dimeric conformation in a similar manner as what Cys3 does, but through a distinct mechanism, i.e. through covalent linkage of the two S100A6 protomers.

## Discussion

In this study, we show that cysteine residues are of critical importance for the binding of S100A6 to the pro-inflammatory RAGE receptor, at least *in vitro*, since removal of S100A6’s unique cysteine at position 3 totally abrogates complex formation with RAGE. This contrasts with the previous report by Mohan *et al* which showed *in vitro* that S100A6-C3S is capable of binding to the RAGE V domain [21]. This discrepancy may be linked to a differential behavior of the isolated RAGE V domain as compared to the full-length ectodomain, with respect to ligand binding. Indeed, the V domain of RAGE is highly basic and may easily interact with acidic proteins in an *in vitro* context, especially in low salt-containing buffers that would favor non-specific electrostatic interactions. Additionally, absence of the C1 and C2 fragments may favor interactions that are otherwise hindered in the complete RAGE protein, highlighting the need to always validate these interaction data on the full-length receptor, and in combination with an evaluation of the calcium-dependency of the interaction for specificity check.

In the structure of the WT RAGE:S100A6 complex, Cys3 from S100A6 is not directly involved in contacts with RAGE [23]. Thus, its major role in promoting S100A6 binding to RAGE is rather linked to its capacity to induce and/or stabilize the RAGE-bound S100A6 dimeric conformation. Our SEC assays show that replacement of Cys3 by a cysteine at conserved position 84 allows to restore full interaction with RAGE. Furthermore, our structure of the RAGE:S100A6-Y84C complex supports the idea that introduction of Cys84 allows to form a RAGE:S100A6 complex identical to the WT one, with a 2:2 stoichiometry and an interaction primarily achieved with the receptor C1 domain. It also demonstrates that Cys84 induces the same RAGE-bound S100A6 homodimeric conformation as Cys3, although through a distinct mechanism involving intermolecular disulfide-crosslinking of the two S100A6 protomers.

While Cys84 is not naturally present in S100A6, a cysteine in the second half of helix H4 is found in at least ten distinct S100 proteins known to bind to RAGE (Fig. 1A), including S100B, S100A2, S100A4, and murine S100A9, that were all reported to form disulfide-crosslinked dimers [15,19,27,28]. Thus, under oxidative conditions, helix H4 cysteines might serve as triggers for the formation of covalent S100 dimers. As a first step towards the corroboration of this hypothesis, modeling of S100A4 and S100B in the RAGE-bound S100A6 dimeric conformation reveals that the conserved cysteine at position 84 (86 for S100A4) would be facing its equivalent residue in the second S100 protomer, the two sulfhydryl groups being within 1.8-2.0 Å distance range, a value compatible with the formation of a disulfide bridge (Figs. 2D-E). Together with Cys68, Cys84 was shown to be important for the formation of S100B covalent dimers [29], which gives strength to our hypothesis. Our structure of the RAGE-bound S100A6-Y84C dimer may therefore represent one possible quaternary organization for these S100 disulfide-crosslinked dimers. The demonstration that such S100 dimeric assemblies exist *in vivo* and the characterization of their precise architecture certainly deserves further investigation, but our data provide preliminary evidence that they can physically exist and that they may be important for RAGE binding.

## Materials and Methods

### Expression and purification of the S100A6 and RAGE proteins (WT and mutants)

The S100A6 C3S, Y84C and C3S-Y84C mutants were generated by PCR-based site-directed mutagenesis from the pETM11:mS100A6 construct previously used to express WT murine S100A6 [23], using anti-complementary primers baring the mutation to introduce and Hot Flex Phusion DNA polymerase (New England Biolabs) according to manufacturer’s instructions. Expression and purification of the recombinant S100A6 proteins was conducted according to previously described procedures [23,30]. Expression and purification of the human RAGE VC1C2 ectodomain (hRAGE-VC1C2; residues Ala23 to Pro323) was done as previously described [31]. Sequence alignments were done in CLUSTAL OMEGA [32] and conservation was analyzed using ALINE [33].

### RAGE:S100 complex formation on size exclusion chromatography (SEC)

SEC binding assays were performed at room temperature on a 24 ml Superdex 75 Increase 10/300 GL column (Cytiva) connected to an Äkta Purifier FPLC system (Cytiva). To assay RAGE:S100A6 complex formation, 600 µg of RAGE VC1C2 were mixed with a 10-fold molar excess of S100A6 (WT or mutants) in 20 mM Tris-HCl pH 7.5, 200 mM NaCl ± 5 mM CaCl_2_, in a final volume of 400 µl. The mixes were incubated for 3 hours at 37°C prior to loading onto the SEC column equilibrated in 20 mM Tris-HCl pH 7.5, 200 mM NaCl. Similar amounts of isolated RAGE-VC1C2 or S100A6 were injected individually on the SEC column, following the same incubation period and conditions, to obtain the elution profiles of the unbound proteins. Fractions of interest from the different elution profiles were analyzed by SDS-PAGE.

### Crystallization of the RAGE:S100A6 mutant complexes

hRAGE-VC1C2 and S100A6-Y84C or S100A6-C3S-Y84C were mixed in a 1:2 molar ratio (RAGE:S100) at a final RAGE concentration of 6 mg/ml in 20 mM Tris-HCl pH 7.5, 200 mM NaCl, 5 mM CaCl_2_. Crystallization experiments were carried out in 96-wells sitting drop plates using commercial screens from Hampton Research and Molecular Dimensions Ltd. Crystals of the RAGE:S100A6-Y84C complex appeared at 4°C over a reservoir containing 0.2 M Zn acetate, 0.1 M Na cacodylate pH 6.5, 9% isopropanol, and with 2 mM of NDSB-221 additive (Hampton Research) in the protein drop. Prior to data collection, the crystals were cryoprotected by soaking into the reservoir solution supplemented with 40% isopropanol followed by flash cooling in liquid nitrogen.

### Data collection, structure determination, model refinement and 3D-modeling

A complete dataset extending to 2.35 Å resolution (Table I) was collected at 100K on the ID30A-1 beamline at ESRF (Grenoble, France). The dataset was processed with XDS [34], revealing a I222 symmetry for the RAGE:S100A6-Y84C crystals with one molecule of RAGE and one molecule of S100A6 per asymmetric unit, like the WT complex. The structure was solved by MR in PHASER [35], using the structure of the WT complex as a search model (PDB_ID 42PY; [23]). Refinement of the model was carried out by alternating cycles of manual rebuilding in Coot [36] and cycles of energy minimization in PHENIX.REFINE [37] using individual ADP refinement and TLS parameterization. Final R_work_ and R_free_ values of respectively 20.43 % and 23.52 % were obtained and the model’s quality was assessed with Molprobity [38]. Atomic coordinates and structure factors were deposited in the Protein Data Bank with accession code 9S2X. Single-omit map generation was done with PHENIX.REFINE using simulated annealing after removing residues 83-85 from S100A6 in the input model. 3D-models for S100B and S100A4 covalent dimers were built manually in Coot from manual superimposition of the protomers on the two subunits of the S100A6-Y84C homodimer. All figures were made with the Pymol Molecular Graphics System, version 0.99rc6, DeLano Scientific LLC.

## Acknowledgements

We thank the beamline staff at the European Synchrotron Research Facility (ESRF, Grenoble, France) for technical assistance during data collection. This work was supported by a grant from the European Community’s H2020 Program Marie-Curie Innovative Training Network INFLANET (https://inflanet.eu/): Grant Agreement nr. 955576. This work was also supported by the French Infrastructure for Integrated Structural Biology (FRISBI, ANR-10-INSB-05-01) for access to the ESRF.

## Author contributions

**Maria Demou:** Investigation, formal analysis (equal), data curation (equal), visualization (equal), writing – original draft preparation (supporting) ; **Laure Yatime:** Conceptualization, methodology, supervision, formal analysis (equal), data curation (equal), visualization (equal), writing – original draft preparation (lead), funding acquisition.

